# Differential Interaction between RAC/ROP-GTPases and RIC-Effectors: A Network Hub for Broader Signal Transduction in Pollen Tubes

**DOI:** 10.1101/2021.04.09.437263

**Authors:** Octavian O. H. Stephan

## Abstract

To date knowledge about plant RAC/ROP-GTPase effectors and downstream targets is still limited. This work aims on elucidation of related signal transduction networks involved in pollen tube growth. Yeast two-hybrid and Pull Down methodology were used to identify and characterize hitherto unknown components of RAC-related protein complexes from *Nicotiana tabacum*. Nt-RIC11pt specifically interacts with diverse active tobacco RAC-GTPases, and it is particularly significant, that their binding affinity is differential, thus implicating a multifaceted role in an interconnected RIC-RAC network. Moreover, Y2H-screening for Nt-RIC11pt targets identified Nt-CAR4, which is phylogenetically assigned to a multifaceted family of novel unusual <u>G</u>TPase <u>a</u>ctivating <u>p</u>roteins (GAP). It is argued that scaffold Nt-RIC11pt connects active Nt-RAC3 with membrane-bound Nt-CAR4, thus relaying GAP-activity. Quantitative RT-PCR demonstrates Nt-RIC11pt is primarily expressed in pollen and YFP-fusion proteins show homogeneous cytoplasmic localization in growing tubes, what builds the prerequisite for a proposed role in broader signal transduction. By synoptically integrating experimental data, bioinformatic sequence comparison, phylogenetic analyses, and detailed literature review, this study hypothesizes a concept in which RIC-effectors collectively constitute a multifaceted network hub linking diverse GTPase-dependent processes.

## 1. INTRODUCTION

Eukaryotic GTP-binding proteins of the Ras superfamily of monomeric small GTPases are essential molecular switches in signal transduction events participating in a variety of physiological processes, such as establishment of cell polarity, cell growth, morphogenesis, and hormone responses (Hakoshima et al., 2003; Kahn et al., 1992; Wennerberg et al., 2005). In species of fungi and animals this Ras superfamily comprises five families, namely RAS, RHO, RAB/YPT, ARF, and RAN, whereas plants have four families as to date no RAS-GTPases could be identified (Kahn et al., 1992; Vernoud et al., 2003; Wennerberg et al., 2005; Winge et al., 1997). The members of each family have been described to be involved in specific cellular events, in particular, RHO-GTPases control actin reorganization, cell polarity, and cell cycle (Boureux et al., 2007; Hodge and Ridley, 2016; Narumiya and Thumkeo, 2018), while RAB and ARF-GTPases regulate distinct steps of vesicular transport and endomembrane protein trafficking (Kjos et al., 2018; Memon, 2004; Pfeffer, 2017). Additionally, the RAN family is related to nucleocytoplasmic transport of RNA and proteins, as well as to mitotic spindle formation (Matsuura, 2016; Zhang and Dawe, 2011).

In contrast to yeast and mammals, which have three subfamilies of RHO-GTPases (Cdc42, Rac, and Rho), plants specifically possess a single subfamily of Rho small GTP-binding proteins termed <u>R</u>ho-like GTPases <u>o</u>f <u>p</u>lants (ROP), also often named as RACs based on their sequence similarities with human Rac-GTPases (Brembu et al., 2006; Feiguelman et al., 2018; Sormo et al., 2006; Winge et al., 2000; Yang, 2002; Zheng and Yang, 2000). Phylogenetic analysis shows that ROPs divided into two distinct evolutionary populations of which one apparently has evolved exclusively in vascular plants (Winge et al., 2000). Moreover, according to sequence similarities all Arabidopsis ROPs are phylogenetically classified into four subgroups with different functions (Gu et al., 2004; Yang, 2002).

Plant RAC/ROP-GTPases are localized in distinct specific subdomains of the plasma membrane and have been identified as central conductors spatially regulating diverse cellular tasks, such as actin cytoskeleton organization and endomembrane transport (Feiguelman et al., 2018; Kost, 2008). RAC/ROPs perform their regulatory function by switching between GTP- and GDP-bound states what facilitates transient interactions with effectors and other regulatory proteins resulting in the control of downstream signalling (Feiguelman et al., 2018; Nagawa et al., 2010). The activity state of small GTPases is spatially regulated by GDP/GTP Exchange Factors (GEFs) that facilitate ROP activation through the exchange of GDP for GTP (‘on’-state), and GTPase-Activating Proteins (GAPs) that enhance the inefficient intrinsic GTP-hydrolysis activity, thereby triggering ROP conformation to the ‘off’-state (Berken and Wittinghofer, 2008; Gu et al., 2006; Moon and Zheng, 2003; Shichrur and Yalovsky, 2006; Wu et al., 2000).

The sequence of some downstream RAC/ROP-binding effector proteins contains a highly conserved CRIB-domain (<u>C</u>dc42/<u>R</u>ac-interactive <u>b</u>inding), which is required for their specific interaction with GTP-bound activated RAC/ROP (Wu et al., 2001).

In addition, CRIB-domains are also found in other RAC/ROP interacting regulatory proteins, such as GAPs or GEFs (Borg et al., 1999; Wu et al., 2000). Whereby the CRIB-domain of classical ROP-GAPs is necessary for both, effective binding to ROP and for enhancing ROP-activity (Klahre and Kost, 2006; Schaefer et al., 2011; Wu et al., 2000).

Several RAC/ROP-effectors have been identified (Nagawa et al., 2010), such as ICR/RIP proteins (Interactor of <u>c</u>onstitutively active <u>R</u>OP/ <u>R</u>OP-interactive <u>P</u>artner)(Hazak et al., 2010; Hazak et al., 2014; Lavy et al., 2007; Li et al., 2008) and different types of RIC proteins (<u>R</u>OP interacting <u>C</u>RIB-motif containing proteins) (Bascom et al., 2019; Gu et al., 2005; Wu et al., 2001). Apart from ICR/RIPs and RICs, some other effector proteins have been identified, such as ABI, RBK, RISAP, and TOR, which are related to ABA-signalling, pathogen response, pollen tube growth, and auxine response (Li et al., 2012a; Li et al., 2012b; Molendijk et al., 2008; Schepetilnikov et al., 2017; Stephan et al., 2014). Interestingly, these types of effectors do not depend on a CRIB-domain to interact with RAC/ROP-GTPases.

ROP-effectors of the ICR/RIP-family in Arabidopsis act as scaffolds, as for instance ICR1/RIP1 (Hazak et al., 2010; Hazak et al., 2014; Lavy et al., 2007; Li et al., 2008), which connects ROPs to the exocyst via SEC3, thus representing an auxin-modulated link between RAC/ROP-signalling, vesicle trafficking, cell polarity, and differentiation.

RICs are a family of proteins with relatively low molecular weight (9 - 25 KDa) that act as direct targets of active ROP-GTPases (Wu et al., 2001). In *Arabidopsis thaliana* 11 RIC genes are known (Wu et al., 2001), and for *Physcomitrella patens* only a single RIC has been described (Bascom et al., 2019), whereas up to now 7 RICs have been predicted by computer analysis of *Nicotiana tabacum* DNA sequences from database. For instance, Arabidopsis ROP1 has been described to bind strongly to RIC1, RIC2, RIC4, RIC5, RIC7, RIC9, and weakly to RIC6 (Wu et al., 2001), whereby RIC4 and RIC3 act as ROP1 effectors in actin-remodelling (Gu et al., 2005). Additionally, RIC1 promotes cortical microtubule organization (Fu et al., 2005) as well as actin filament severing (Zhou et al., 2015).

However, Arabidopsis At-RIC11 has not been characterized, and Wu et al. (2001) state that isolation of this gene turned out to be problematic (Wu et al., 2001). Thus, the present work provides first information for the RIC11-type of effector proteins, showing that *Nicotiana tabacum* RIC11 from pollen tubes is a true interactor of active RAC/ROP-GTPases from different phylogenetic groups representing diverse cellular pathways.

Moreover, observations provided in the present study demonstrate that Nt-RIC11pt binds to Nt-CAR4 which is a homolog of At-GAP1 and Os-GAP1, thus it is concluded that effectors of the RIC-family do not only transmit signals to downstream targets, but Nt-RIC11pt for its part also relays feedback to RAC/ROPs by recruiting a special type of GTPase-Activating Protein (GAP) family.

By integrating literature study with primary research the goal of this work is to contribute novel insights into the character of plant RAC/ROP-effectors by providing a hypothesis how RICs are collectively involved in signal transduction during polar cell growth exemplary of *Nicotiana tabacum* pollen tubes.

While the eclectic Rho-GTPase family of humans consists of 20 members (Jaffe and Hall, 2005; Narumiya and Thumkeo, 2018; Wennerberg et al., 2005), the unique ROP-family of plants on the other hand comprises substantially less members, as for instance 4 ROPs in *Physcomitrella patens*, 7 in *Oryza sativa*, and 11 in Arabidopsis (Brembu et al., 2006; Feiguelman et al., 2018; Li et al., 1998). Data describing how these molecular switches take effect are still limited and only some members of downstream signal transduction chains have been characterized in plants. In comparison to humans the obviously lower number of Rho-GTPases in plants raises the intriguing question how this restricted quantity is able to regulate the multitude of related processes. A potential plant specific answer to this particularly important question might be a multifaceted character of RIC-effectors interacting with various GTPases. In that regard, by integrating the observation that Nt-RIC11pt binds with differential affinity to diverse RAC/ROP-GTPases with data from several other studies the present article aims on demonstrating that the RIC-type of effectors represent a molecular network hub interconnecting various signalling pathways.

## 2. MATERIALS AND METHODS

### 2.1 Yeast cultivation and two-hybrid screen

Yeast strains were cultivated on YEP-based media with 2% glucose (YEPD) or synthetic complete (SCD) plates supplemented with amino acids in an incubator for 3-4 days at 30° (Stephan and Koch, 2009). All yeast two-hybrid screens and assays were performed using the BD Matchmaker system (BD Biosciences, Clontech Laboratories) utilizing the *Saccharomyces cerevisiae* strain AH109 essentially as described earlier (Stephan et al., 2014). Plasmid constructs based on vectors pGBKT7 (bait) and pGADGH (prey) were co-transformed into yeast and transformants were selected on SCD-medium lacking tryptophan, leucine, and histidine.

### 2.2 Plant cultivation, pollen tube transformation and microscopic analysis

*Nicotiana tabacum* (*cv Petite Havana SR1*) plants were cultivated on soil under standard conditions (Stephan et al., 2014). Pollen were germinated and pollen tubes cultured at 25°C in liquid PTNT medium (Read et al., 1993a; Read et al., 1993b). For transient pollen tube transformation the pollen was released from anthers and transferred on solid PTNT medium plates before biolistic particle bombardment with DNA-covered gold particles as described (Stephan et al., 2014). Transformed tobacco pollen tubes were grown for six hours at 25°C on PTNT plates. Thereafter a piece of solid medium together with growing pollen tubes was cut out and transferred upside-down onto cover slips. Confocal images were generated using a Leica TCS SP2 laser scanning microscope (PLAN FLUAR 1003/1.45 oil immersion objective; a HC PL APO 203/10.7 water immersion objective). YFP fluorescence was excited at 514 nm and detected between 530 and 600 nm.

### 2.3 RNA analysis and real time qPCR

Pollen, pollen tubes, and fresh tobacco tissue samples were frozen in liquid nitrogen. Tissue cells were disrupted and homogenized by a tissue lyser in a denaturing buffer containing guanidine-salts. Total RNA was isolated using the RNeasy Plant Mini Kit according to the manufacturer’s protocol (QIAGEN, Germany) and eluted in 30μl nuclease free water. Total RNA was treated with DNAse before undergoing reverse transcription. For analysis of isolated RNA semi-quantitative reverse transcription–polymerase chain reaction (RT) PCR was performed with the iScript™ cDNA Synthesis Kit (BIORAD, Germany) according to the manufacturer’s instructions using 500ng of isolated RNA. Primers specific to Nt-RIC11 and Nt-LF25 (Accession number: L18908) were used in multiplex semi-quantitative PCR, amplifying products of 260 and 380 bp. Quantitative cDNA detection was performed via quantitative real-time PCR (RT-qPCR) with the same gene-specific primers using the iQ™ SYBR^®^ Green Supermix Kit (BIORAD). Serially diluted genomic DNA was used as a quantification standard.

### 2.4 Plasmid construction and cloning of RAC/ROP-GTPases

DNA manipulations, cloning and analysis were carried out according to standard techniques (Sambrook and Russell, 2001). Plasmid constructs for pull down were generated by amplifying coding sequences of target RAC genes with gene-specific primers from cDNA that was reverse-transcribed from isolated tobacco pollen tube RNA. The gene-specific forward and backward primers were designed in accordance with the 5‘and 3’ends (from Start to Stop codon) of existing and predicted tobacco RAC DNA-sequences (see chapter 2.7 Accession numbers) which were retrieved from Genebank database (https://www.ncbi.nlm.nih.gov/gene). EcoRI and XhoI restriction sites, which were added to the primer sequences, allowed to clone the complete amplified coding sequence of the respective tobacco RAC in pGADGH and pGEX-4T2 vectors. The obtained cDNA clones were verified by DNA-sequencing and BLAST searches.

### 2.5 Pull down assays

GST-fusion proteins were expressed in *E. coli* BL21 (DE3) cells that were transformed with constructs derived from pGEX-4T-2 vector containing different cDNA inserts. Purification of GST-fusion proteins from cell extracts and nucleotide loading of RAC/ROP-GTPases was performed as described earlier (Stephan et al., 2014). Pull down assays were performed utilizing myc-tagged proteins that were generated from recombinant pGBKT7-derivatives through *in vitro* transcription/translation using wheat germ extract (cat# L4330, Promega, Madison, WI, USA)(Stephan et al., 2014).

### 2.6 Software tools

Sequence analysis of proteins was performed with following software tools: Dompred (http://bioinf.cs.ucl.ac.uk/psipred/?dompred=1); PROSITE (https://prosite.expasy.org); GPS3.0 software (http://gps.biocuckoo.org/). Sequence percent identity matrix was created by Clustal2.1 software. Phylogenetic trees were generated using the ‘MEGA-X 10.0.5’– software (https://www.megasoftware.net/).

### 2.7 Accession numbers

DNA, RNA and Protein Sequence data are documented in the GenBank/EMBL database under following accession numbers:

**Nt-RAC1 = MN786790** *[similar to XP_016464941 / XM_016609455 / LOC107787847 (identical to protein AAK31299, but this has 19 nucleotide changes in mRNA sequence AY029330 / LOC107768360)*].

**Nt-RAC3 = MN786791** [*similar to XP_016469644 / XM_016614158 / LOC107791990*].

**Nt-RAC5 =** *XP_016476800 / AJ222545 / LOC107798337; XP_016512160 / XP_016512162 / XP_016512161 / AJ250174*.*1 / LOC107829207; AAD00117 (All proteins are identical, but have different mRNA sequences)*.

**Nt-RAC5.2 = MN786792** [*similar to Nt-RAC5, but mRNA has 18 nucleotide changes and protein has one P to S substitution*].

## 3. RESULTS

### 3.1 Identification of tobacco RIC11 from pollen tubes using yeast two-hybrid

To date only two RAC/ROP-GTPases have been described in *Nicotiana tabacum*, namely Nt-RAC1 (Tao et al., 2002) and Nt-RAC5 (Kieffer et al., 2000), whereby the most comprehensively analysed is Nt-RAC5 (AJ250174)(Kieffer et al., 2000; Klahre et al., 2006; Stephan et al., 2014). Additionally, the tobacco NTGP2 (AAD00117) protein has to be mentioned which is identical to Nt-RAC5, however, their mRNA sequences differ.

Nt-RAC5 has been related to the control of polarized pollen tube growth (Kieffer et al., 2000) thereby regulating actin dynamics and membrane transport (Stephan et al., 2014).

To contribute novel information to the Nt-RAC5 effector network yeast two-hybrid (Y2H) screening methodology was used to identify proteins interacting with a constitutive active tobacco RAC5 mutant [G15V], because many downstream effector proteins, such as particularly CRIB-domain containing proteins, have been described to bind Rho in the GTP-bound “active” state (Feiguelman et al., 2018; Nagawa et al., 2010). In this screen of a pollen tube library several partial cDNAs of unknown proteins were identified as potential Nt-RAC5 downstream effectors, and in addition, a cDNA encoding an N-terminal truncated fragment (amino acids 9-144) of a protein belonging to the family of CRIB-motif containing proteins (**Fig.1A**). Cloning of the corresponding full-length cDNA was performed by a PCR based strategy using gene specific primers, resulting in an open reading frame encoding a 144-amino acid (15-kD) protein (**Fig.1A,B**). BLAST search for Arabidopsis homologs revealed that this protein is most closely related to At-RIC11 (AT4G21745; 38% sequence identity) and At-RIC10 (AT4G04900.1; 36.3% identity) (**Fig.2A**).

**Figure 1.**
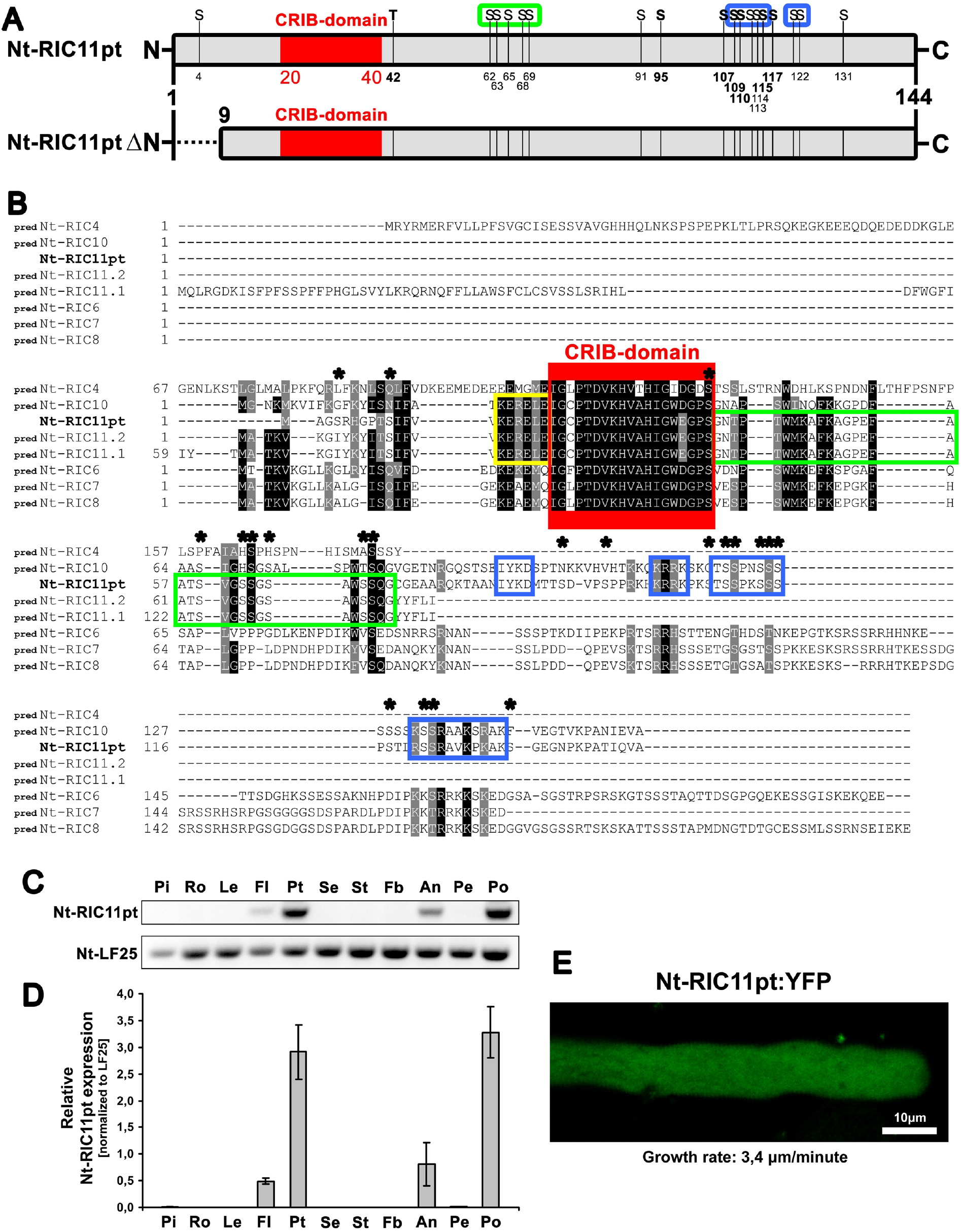
Domain Structure, Amino Acid Sequence Comparison, Tissue Specific Expression Levels, and Subcellular Localization of Tobacco Nt-RIC11pt. **(A)** Domain structure of full-length Nt-RIC11pt protein (aa 1-144; top) and the N-terminal truncated version (aa 9-144) identified in the yeast two-hybrid screen as Nt-RAC5 interaction partner. S highlight the position of serine residues in Nt-RIC11pt predicted as potential phosphorylation sites via GPS3.0 software; Bold S and T highlight the serine and threonine residues consistently predicted by GPS3.0, NetPhos 3.1 (http://www.cbs.dtu.dk/services/NetPhos/), and KinasePhos (http://kinasephos.mbc.nctu.edu.tw/) software. Red box: CRIB-motif; Blue boxes: Serine residues homologous between Nt-RIC11pt and predicted Nt-RIC10(XP_016467450); Green box: Serine residues conserved between _**pred**_Nt-RIC11pt and _**pred**_Nt-RIC11 isoforms Nt-RIC11.1(XP_016507046) and Nt-RIC11.2(XP_016507047). **(B)** Clustal Omega Amino Acid Sequence Alignment (https://www.ebi.ac.uk/Tools/msa/clustalo/) of Nt-RIC11pt with 7 predicted *Nicotiana tabacum* RICs identified by BLAST search. The tobacco _**pred**_Nt-RIC sequences were predicted by automated computational analysis of genomic sequences (HMM-based gene prediction program *Gnomon* (https://www.ncbi.nlm.nih.gov/genome/annotation_euk/gnomon/). Black shading: sequence identity; Gray shading: sequence similarity; Red box: CRIB-motif; Blue boxes: Amino acid sequences specifically conserved between isolated Nt-RIC11pt and _**pred**_Nt-RIC10 (XP_016467450); Green boxes: Amino acid sequences specifically conserved between isolated Nt-RIC11 and predicted _**pred**_Nt-RIC11 isoforms _**pred**_Nt-RIC11.1 (XP_016507046) and _**pred**_Nt-RIC11.2 (XP_016507047); Yellow box: Amino acids conserved between Nt-RIC11pt, predicted _**pred**_Nt-RIC10 and both predicted _**pred**_Nt-RIC11 isoforms; Black asterisks mark all serine residues in Nt-RIC11pt; Nt: *Nicotiana tabacum*. **(C)** Semi-quantitative RT-PCR analysis of Nt-RIC11t mRNA levels in different wild-type tobacco tissues compared to mRNA levels of the Nt-LF25 reference gene. The mRNA was purified from different plant tissues and used as template for cDNA synthesis. Corresponding cDNA was analysed with gene specific primers via multiplex PCR and the amplified products were stained with ethidium bromide and analysed on a 2% agarose gel. Pi: pistils; Ro: roots; Le: leaves; Fl: flowers; Pt: pollen tubes; Se: sepals; St: stems; Fb: flower buds; An: anthers; Pe: petals; Po: pollen. **(D)** Nt-RIC11pt mRNA levels in different wild-type tobacco tissues analysed by quantitative reverse transcription real time polymerase chain reaction (qRT-PCR) assay using gene specific primers for Nt-RIC11pt and the Nt-LF25 reference gene (Materials and Methods). Displayed is the mean relative Nt-RIC11pt mRNA level of three biological replicates normalized to Nt-LF25 mRNA. Pi: pistils; Ro: roots; Le: Leaves; Fl: flowers; Pt: pollen tubes; Se: sepals; St: stems; Fb: flower buds; An: anthers; Pe: petals; Po: pollen. **(E)** Single confocal optical section through a normal growing pollen tube transiently expressing Nt-RIC11pt:YFP 6h after gene transfer. The Nt-RIC11pt fusion protein shows a clear cytoplasmic distribution pattern. Bar = 10 μm.

To this day no protein sequences of *Nicotiana tabacum* RIC-family members were experimentally verified, however, computer predictions exist in NCBI database, whereby predicted _pred_Nt-RIC10 shows high sequence similarity (64%) while _pred_Nt-RIC11 has the best match (84.2%) (**Fig.1B**), therefore the isolated ORF from pollen tubes is called Nt-RIC11pt hereafter.

Further homologs were identified in several other plant species, with nearest in Solanaceae, Asteraceae, and Salicaceae, while outside of plant kingdom only two were identified, namely the closest non plant homolog *Homo sapiens* WASP (Kim et al., 2000)(22.4% identity), and *Saccharomyces cerevisiae* STE20 (15.4%). Interestingly, Sc-STE20 is involved in polarized pseudohyphal growth of yeast and the pheromone response pathway, whereby Sc-STE20 is a direct interaction partner of the yeast CDC42 Rho-like GTPase (Bardwell, 2005; Gancedo, 2001) which is the homolog of tobacco Nt-RAC5. Additionally, the human Wiskott–Aldrich syndrome protein (WASP), which contains a CRIB-domain, has been identified as a direct effector of Hs-CDC42 regulating actin-dynamics (Abdul-Manan et al., 1999; Kim et al., 2000). Thus, Nt-RIC11pt, Sc-STE20, and Hs-WASP correspondingly interact with CDC42 related small GTPases, what allows to hypothesize an effector function for tobacco Nt-RIC11pt in polarized growth and actin-dynamics, as well as a potential role in intracellular transduction of extracellular signals.

### 3.2 Protein structure of Nt-RIC11pt

Interestingly, the experimentally verified Nt-RIC11pt does not fully match the predictions and thus appears to be a hybrid-like protein combining the computed sequence characteristics of both, _pred_Nt-RIC10 and _pred_Nt-RIC11 (**Fig.1B**). Sequence alignments with all predicted tobacco RICs demonstrate that the isolated Nt-RIC11pt shares some amino acids exclusively either with _pred_Nt-RIC10 (**Fig.1B**; blue boxes), or _pred_Nt-RIC11 (**Fig.1B**; green box), whereas the CRIB-domain and an adjoining region are highly conserved (**Fig.1B**; red box; yellow box).

A striking feature is the exceptional number of clustered serine residues in Nt-RIC11pt (**Fig.1B**; asterisks), which primarily accumulate C-terminally of the CRIB-domain in areas either similar to _pred_Nt-RIC10 (**Fig.1A**; blue boxes) or _pred_Nt-RIC11 (**Fig.1A**; green box). Through sequence analysis with GPS3.0, NetPhos 3.1, and KinasePhos software several putative phoshorylation sites were identified (**Fig.1A**; bold S and T).

Based on alignments including all known Arabidopsis and predicted *Nicotiana tabacum* RIC proteins a phylogenetic tree was generated using maximum likelihood estimation with 500 replications (**Fig.2C**). The unrooted tree and amino acid similarities resulted in categorization of all RICs in five groups, whereby Nt-RIC11pt belongs to group I, comprising RIC9, RIC10, and RIC11 (**Fig.2C**). Sequence alignment of tobacco Nt-RIC11pt with Arabidopsis group I members indicates that amino acid similarity is limited to the CRIB-domain, a conserved consensus sequence and several serine residues (**Fig.2A**). Interestingly, sequence similarity of the complete RIC-family is significantly low among its members (**Fig.2B**) and primarily restricted to the highly conserved CRIB-domain, a conserved consensus sequence (KX[RK]N[KR]KXK) and an adjacent conserved serine/threonine site (**Fig.3A,B**). The localization of these conserved structural features within the amino acid sequence of RICs evidently specifies members of the respective phylogenetic group (**Fig.3B**).

**Figure 2.**
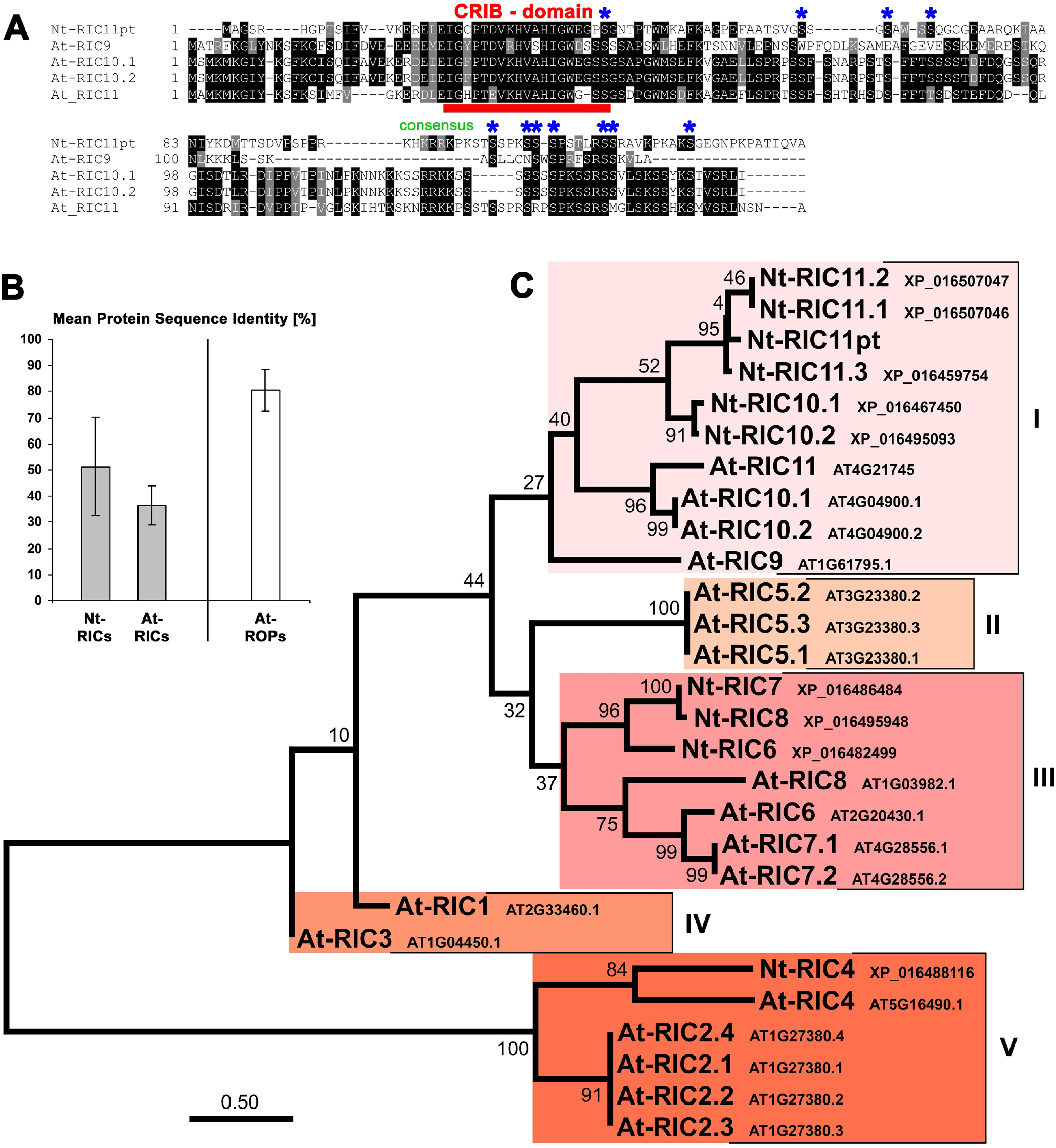
Comparison of Nt-RIC11pt with Closest Homologs from *Arabidopsis thaliana* and predicted *Nicotiana tabacum* RICs . **(A)** Clustal Omega Amino Acid Sequence Alignment of Nt-RIC11pt with *Arabidopsis thaliana* At-RIC9 (AT1G61795.1), At-RIC10.1 (AT4G04900.1), At-RIC10.2 (AT4G04900.2), and At-RIC11 (AT4G21745) from TAIR database. Black shading: sequence identity; Gray shading: sequence similarity; Blue asterisks mark all serine residues conserved between Nt-RIC11pt, At-RIC9, At-RIC10, and At-RIC11; Red bar: highly conserved CRIB domain; Nt: *Nicotiana tabacum*; At: *Arabidopsis thaliana*. **(B)** Statistical analysis of protein sequence identities of 18 known Arabidopsis RICs, 9 computer predicted tobacco RICs, and for comparison of 11 known Arabidopsis ROPs. Indicated are their respective mean sequence identities [%] and the standard deviation. **(C)** Phylogenetic analysis delineated in an unrooted tree obtained by maximum likelihood estimation and JTT matrix-based model providing protein amino acid sequence relationships between Nt-RIC11pt, known At-RICs from *Arabidopsis thaliana*, and computer predicted Nt-RICs of *Nicotiana tabacum* from database search. All RICs were classified in V groups (various red tones) according to sequence similarity. Phylogeny reconstruction was generated using ‘MEGA-X 10.0.5 - software’. The indicated bootstrap values are based on 500 replications and the scale bar for branch lengths indicates the number of amino acid substitutions per site. Locus/Accession number is given for every protein. At: *Arabidopsis thaliana*; Nt: *Nicotiana tabacum*.

**Figure 3.**
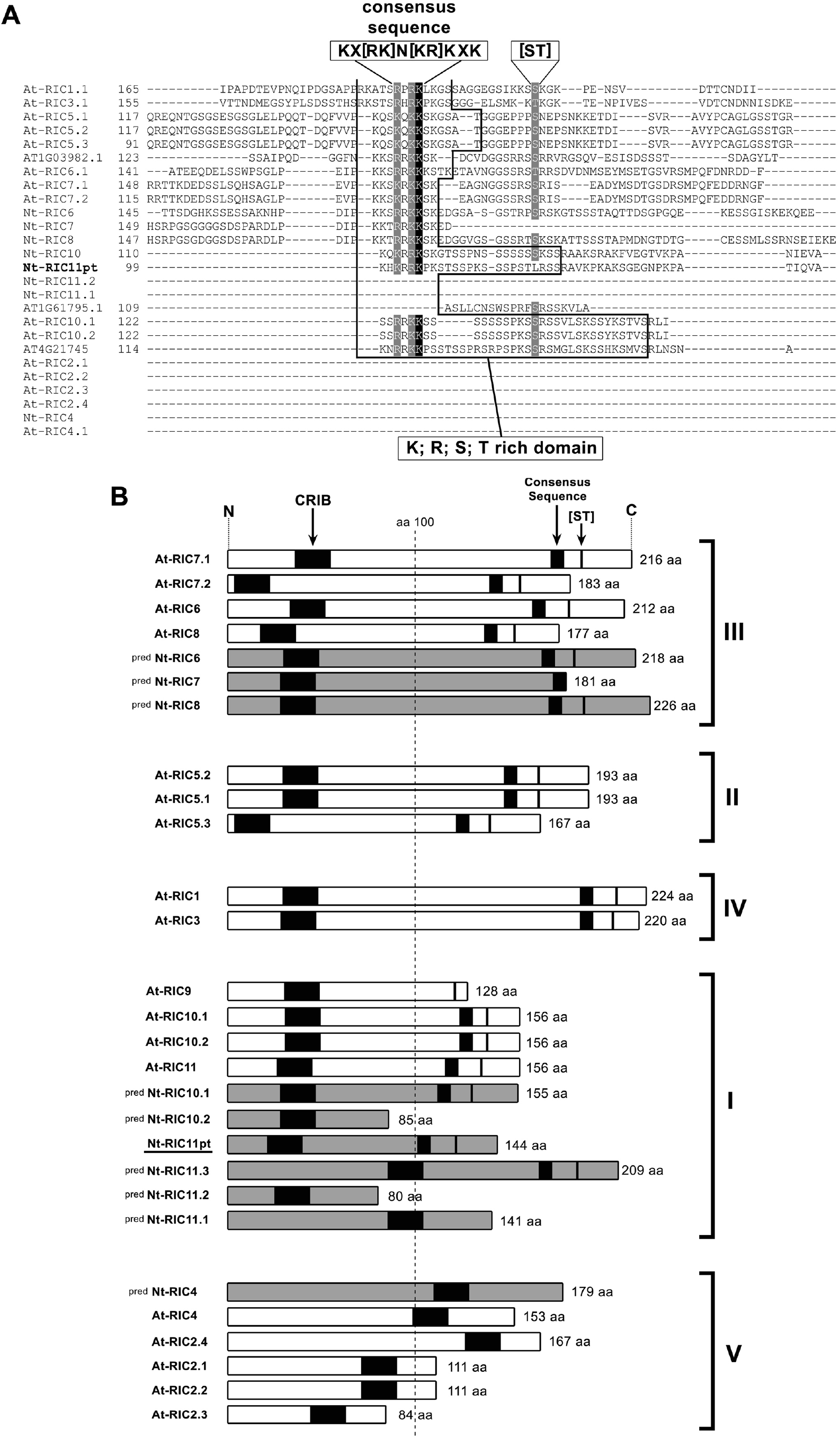
Domain Structure Comparison of all known Arabidopsis and predicted Tobacco RICs. **(A)** Clustal Omega Amino Acid Sequence Alignment of Nt-RIC11pt with all known Arabidopsis homologs and 7 predicted *Nicotiana tabacum* RICs identified by database search. The tobacco _pred_Nt-RIC Sequences were predicted by automated computational analysis of genomic sequences (HMM-based gene prediction program *Gnomon* (https://www.ncbi.nlm.nih.gov/genome/annotation_euk/gnomon/). Indicated are the highly conserved consensus sequence and the conserved serine/threonine [ST] residue. Black shading: sequence identity; Gray shading: sequence similarity; At: *Arabidopsis thaliana*; Nt: *Nicotiana tabacum*. **(B)** Protein domain structure comparison of Nt-RIC11pt with all known Arabidopsis At-RICs (white) and 7 predicted *Nicotiana tabacum* RICs (grey) identified by database search. RICs were classified in V groups according to sequence similarity. Black boxes: indicate the CRIB-domain, as well as the consensus sequence and the conserved serine/threonine residue in the K/R/S/T rich domain; pred.: predicted; aa: amino acid; At: *Arabidopsis thaliana*; Nt: *Nicotiana tabacum*.

#### 3.3 Tissue-dependent expression profile and subcellular localization of Nt-RIC11pt

To identify the tissue-dependent expression pattern of Nt-RIC11pt its transcript levels were investigated from total RNA samples obtained from various tobacco plant organs. Semi-quantitative RT-PCR analysis indicates that Nt-RIC11pt mRNA accumulates predominantly to high levels in tobacco pollen and pollen tubes (**Fig.1C**). Additionally Nt-RIC11pt transcripts were also found at low levels in anthers and even smaller quantities in mature flowers, what is most probably due to the fact that pollen are components of these tissue samples.

These results were confirmed by quantitative real-time PCR analysis of derived cDNA from three biological replicates (**Fig.1D**). The quantitative data of Nt-RIC11pt transcript levels were normalized to mRNA levels of reference genes, such as the constitutively expressed tobacco LF25 ribosomal protein (**Fig.1D**). Interestingly, this distinct expression pattern, showing highest Nt-RIC11pt levels in pollen and pollen tubes, correlates with the one described for tobacco Nt-RAC5 (Klahre et al., 2006) and Arabidopsis At-ROP1, additionally, At-ROP3, At-ROP5, At-ROP11 show expression profile overlaps in pollen and flowers, but are also allocated to other vegetative tissues (Dorjgotov et al., 2009; Li et al., 1998).

To gain insight into the subcellular localization of Nt-RIC11pt in pollen tubes, YFP (yellow fluorescent protein) was fused to the C-terminus of Nt-RIC11pt (Nt-RIC11pt:YFP) and plasmids encoding the constructs were transiently transformed in tobacco pollen by biolistic particle bombardment. Six hours after gene transfer normal growing pollen tubes transiently expressing Nt-RIC11pt:YFP under control of the LAT52 promoter were analysed by confocal microscopy to determine the intracellular distribution. Nt-RIC11pt:YFP was detected evenly distributed in the pollen tube cytoplasm (**Fig.1E**), what resembles the reported localization pattern of related Arabidopsis RIC10 (Wu et al., 2001) from the same phylogenetic group (**Fig.2C**). Altogether, these data, which were derived from *in vitro* cultured pollen tubes, indicate that Nt-RIC11pt is predominantly expressed at high levels in mature pollen and in growing pollen tubes, particularly localized in the cytoplasm, what suggests that Nt-RIC11pt may play a role during pollen germination and tube growth.

### 3.4 Interaction of Nt-RIC11pt with Nt-RAC5 in yeast two-hybrid and pull down assay

The highly conserved CRIB-domain of CDC42/RAC-effectors binds to GTPases in a GTP-dependent manner. To investigate if Nt-RIC11pt interacts specifically with the GTP-bound active form of RAC5-GTPase yeast-two hybrid assays were performed with constitutively active RAC5[G15V], dominant negative RAC5[T20N], and wild-type RAC5 fused to the DNA-binding domain of GAL4 transcription factor (GAL4-DBD). Growth of yeast transformants demonstrates that full-length tobacco Nt-RIC11pt fused to the activation domain of GAL4 (GAL4-AD:Nt-RIC11pt) interacted with ca RAC5[G15V], and to a lesser extent with wild-type RAC5, however, no two-hybrid interaction was detected with dn RAC5[T20N] (**Fig.4A**), which is an inactive mutant form of RAC5 representing the conformation of its nucleotide-free transition state (Feig, 1999; Stephan et al., 2014).

In pull down assays, heterologously expressed N-terminally GST-tagged Nt-RIC11pt was purified from *E. coli* via glutathione-coupled bead matrix. The immobilized GST:Nt-RIC11pt fusion interacted specifically with myc-tagged wild-type RAC5 (myc:Nt-RAC5), ca RAC5[G15V], and dn RAC5[T20N], which were produced by *in vitro* transcription/translation (**Fig.4B**). Immobilized GST alone served as control for specificity of binding. Immunological detection of epitope-tagged fusion proteins with anti-myc and anti-GST antibodies in the pull down precipitates demonstrates a significant interaction specifically between GST:Nt-RIC11pt and myc-tagged ca RAC5[G15V] (**Fig.4B**), whereas on the other hand, the reduced binding to wild-type RAC5 and equally to dn RAC5[T20N] suggests an additional GTP-independent constitutive interaction between Nt-RIC11pt and Nt-RAC5 on a weaker basal level (**Fig.4B**). Most importantly, **figure 4C** shows that this weak constitutive binding of Nt-RIC11pt to wild-type Nt-RAC5 in its inactive GDP-loaded conformation becomes significantly more pronounced when wild-type Nt-RAC5 is in the GTP-loaded active state (**Fig.4C**).

**Figure 4.**
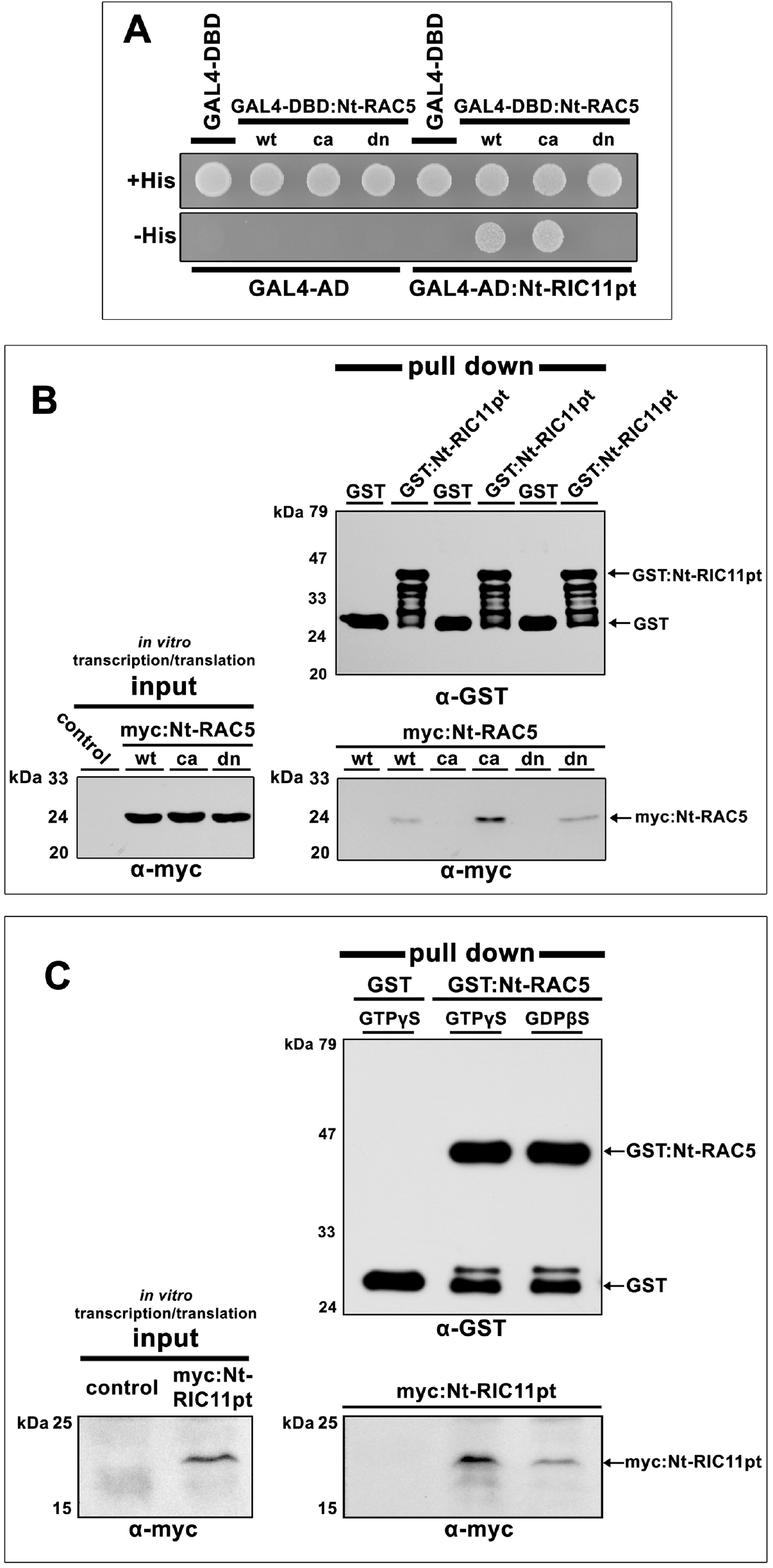
Interaction of Nt-RIC11pt with Constitutively Active, Dominant Negative as well as GDP- or GTP-Loaded Wild-type forms of RAC5 in Pull Down Assays and Yeast Two-Hybrid Assay. **(A)** Yeast transformants co-expressing either wild-type (wt), or constitutive active (ca: G15V), or dominant-negative (dn: T20N) Nt-RAC5 fused to the DNA binding domain of the GAL4 transcription factor (GAL4-DBD) together with Nt-RIC11pt fused to the GAL4 activation domain (GAL4-AD). Cells from single yeast transformant colonies were diluted in water and equal amounts plated on histidine-free culture medium (-His) to analyse for two-hybrid interaction. As positive growth control all transformants were also plated on histidine containing medium (+His). Controls for specific protein interaction were transformants co-expressing the Nt-RIC11pt prey protein with just the GAL4-DBD, as well as transformants co-expressing Nt-RAC5 bait protein together with just the GAL4-AD. **(B)** Equal amounts of GST-tagged Nt-RIC11pt protein were immobilized on glutathione-coupled bead matrix, washed and purified from E. coli cell extracts. Matrix-bound GST:Nt-RIC11pt fusion protein was incubated with *in vitro* transcribed/translated myc:Nt-RAC5 (either wild type [wt], or constitutive active [ca], or dominant-negative [dn] versions of Nt-RAC5). Beads loaded with GST served as control. Precipitated GST:Nt-RIC11pt and co-purified myc-tagged Nt-RAC5 proteins were separated via SDS-PAGE and analysed by immunoblotting applying anti-GST antibodies (upper panel) and anti-myc antibodies (lower panel). Input panel on the left shows in vitro transcription/translation of myc:Nt-RAC5 fusion proteins, which were subsequently subjected to pull down assay. GST: Glutathione-S-Transferase affinity tag; Nt: *Nicotiana tabacum*. **(C)** GST-tagged wild type Nt-RAC5 was purified from E. coli cell extracts via magnetic beads. The matrix-bound GST:Nt-RAC5 fusion proteins were either loaded with GDPβS or GTPγS and then incubated with in vitro-transcribed/translated myc:Nt-RIC11pt. Beads loaded with GST served as control. Precipitated GST-tagged Nt-RAC5 and co-purified myc-tagged Nt-RIC11pt proteins were separated via SDS-PAGE and analysed by immunoblotting applying anti-GST antibodies (upper panel) and anti-myc antibodies (lower panel).

In summary, data of **figure 4** demonstrate that Nt-RIC11pt is an interactor of GTP-loaded active Nt-RAC5, however, also a constitutive GTP-independent binding on a basal comparably lower level was additionally shown. In this regard, Y2H detects an interaction of Nt-RIC11pt with constitutively active and wild-type RAC5, but not with dominant negative mutant RAC5, whereas Pull down assays, on the other hand, present a more detailed picture by clearly indicating a basal interaction between Nt-RIC11pt and wild-type Nt-RAC5, which becomes significantly more distinct in a GTP-dependent manner.

### 3.5 Nt-RIC11pt interacts with different types of RAC/ROP-GTPases

Arabidopsis ROP1 has been reported to interact with different types of At-RICs, such as At-RIC1, At-RIC2, At-RIC4, At-RIC5, At-RIC7, and At-RIC9 (Wu et al., 2001), however, they did not analyse a potential interaction between At-ROP1 and At-RIC11. Considering the existing data of Wu et al. (Wu et al., 2001) their observations would mean by implication that Nt-RIC11pt may potentially bind to various tobacco RAC/ROP GTPases. To check if tobacco Nt-RIC11pt is limited to interaction exclusively with Nt-RAC5, and to examine if Nt-RIC11pt is involved in additional pathways regulated by other RAC/ROP-GTPases, the physical interaction with Nt-RAC1, Nt-RAC3, Nt-RAC5, and Nt-RAC5.2 has been analysed by comparison. Therefore, pull down assays were performed, in which these tobacco RACs were heterologously expressed as N-terminal GST-tagged fusion proteins in *E. coli*, allowing their immobilization and purification via affinity-matrix. The RAC-DNA sequences were isolated with gene-specific primers via PCR amplification of cDNA, which was reverse transcribed from tobacco pollen tube RNA (Materials and Methods). The translated protein sequence of isolated Nt-RAC3 cDNA (MN786791) is identical with that of computer predicted tobacco Rac-like GTP-binding protein 3 (LOC107791990) and shows highest sequence homology with Arabidopsis At-ROP11 (86.6% identity) and At-ROP10 (86.3% identity) (**Fig.5D**). The experimentally obtained Nt-RAC1 (MN786790) is identical with tobacco RAC1 protein (**Fig.5D**) (see chapter 2.7 Accession numbers; AAK31299) (Chen et al., 2003; Tao et al., 2002) and has highest sequence homology with At-ROP1 (96.95% identity) and the related group members At-ROP3 (96.4% identity) and At-ROP5 (95.4% identity) (**Fig.5D**). The isolated Nt-RAC5.2 represents an isoform 351 of Nt-RAC5 (LOC107829207; LOC107798337; AAD00117) (Kieffer et al., 2000; Stephan et al., 2014). Pull Down experiments demonstrate that *in vitro* transcribed/translated myc:Nt-RIC11pt significantly interacted specific with GTPγS-loaded GST-tagged Nt-RAC1, Nt-RAC3, Nt-RAC5, and Nt-RAC5.2 (**Fig.5A**).

Pull down assays were performed in quadruplicate to control for variations in the efficiency of precipitation. As an additional control binding of myc:Nt-RIC11pt to GST alone was analysed in parallel. No significant interaction was detected between GST and myc:Nt-RIC11pt (**Fig. 5A**). Interestingly, digital quantification of the western blot signals demonstrates that equal quantities of myc:Nt-RIC11pt were co-precipitated proportionally with GST:Nt-RAC1, GST:Nt-RAC5, and GST:Nt-RAC5.2, indicating that the interactions of Nt-RIC11pt with these three GTPases are equally strong (**Fig.5B**). On the other hand, almost twice as much Nt-RIC11pt has been co-precipitated proportionally with Nt-RAC3, what suggests that Nt-RIC11pt has a much higher affinity to Nt-RAC3 compared to Nt-RAC1, Nt-RAC5, and Nt-RAC5.2 (**Fig.5B**).

**Figure 5.**
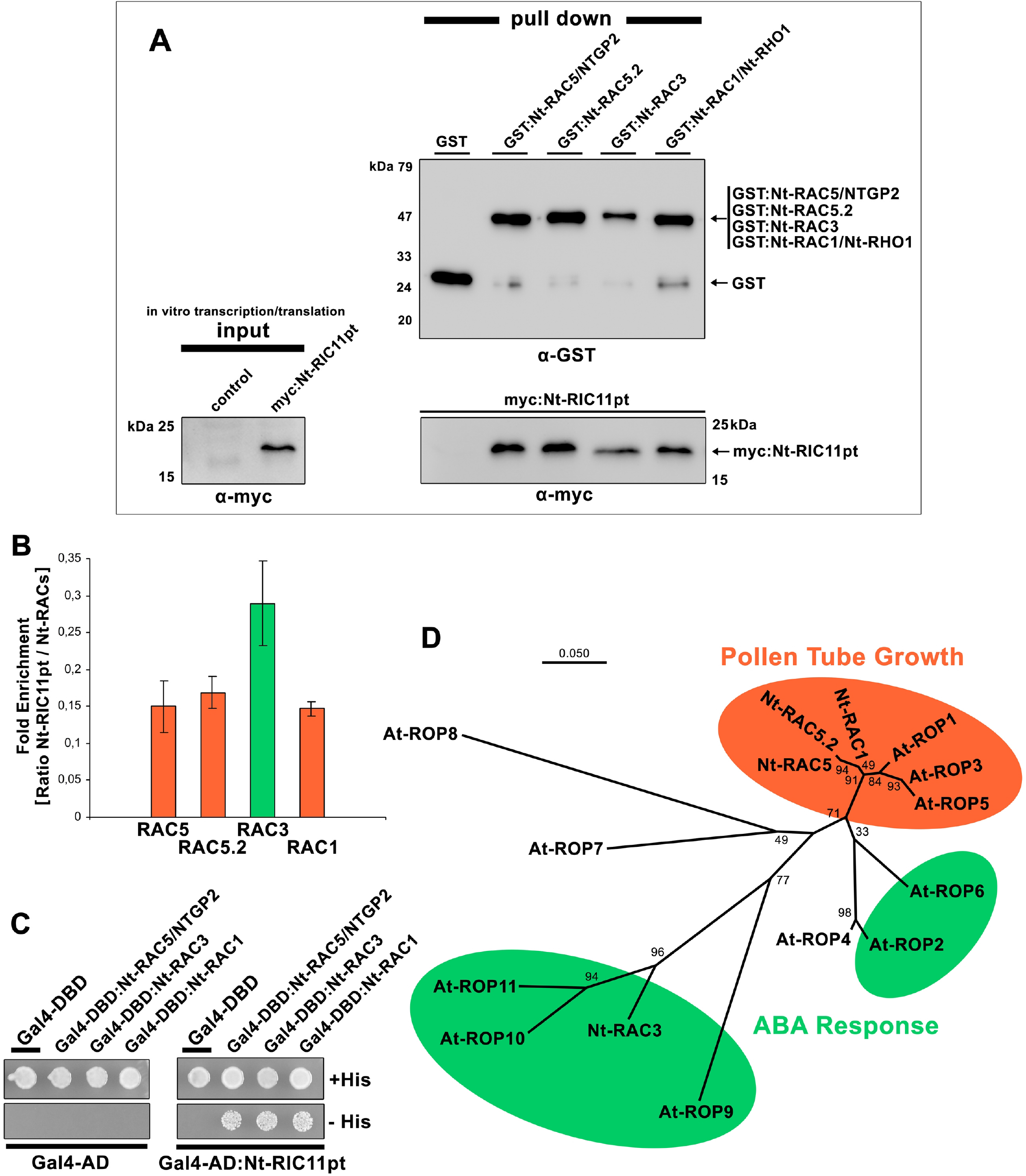
Interaction of Nt-RIC11pt with Nt-RAC3, Nt-RAC5, Nt-RAC5.2, and Nt-RAC1, in Pull Down and Yeast Two-Hybrid Assays. **(A)** GST-tagged Nt-RAC proteins (RAC1, RAC3, RAC5, RAC5.2) were immobilized on magnetic bead matrix, washed and purified from E. coli cell extracts. Each matrix-bound GST:Nt-RAC fusion protein was equally loaded with GTPγS and incubated with *in vitro*-transcribed/translated myc:Nt-RIC11pt. Beads loaded with GST served as control. Precipitated GST-tagged Nt-RACs and co-purified myc-tagged Nt-RIC11pt proteins were separated via SDS-PAGE and analysed by immunoblotting applying anti-GST antibodies (upper panel) and anti-myc antibodies (lower panel). Input panel on the left shows *in vitro* transcription/translation of myc:Nt-RIC11pt fusion proteins, which were subsequently subjected to pull down assay. GST: Glutathione-S-Transferase affinity tag. **(B)** Statistical analysis of protein quantities purified in pull down experiment from (A). Displayed are mean values of the ratio representing myc:Nt-RIC11pt enrichment relative to precipitated GST:Nt-RAC. Co-precipitated myc:Nt-RIC11pt quantities were individually normalized to the myc:Nt-RIC11pt co-precipitate of the GST-background control beforehand. The signal intensities of chemoluminescence detection were determined by digital imaging and quantified by ImageJ software (https://imagej.nih.gov/ij/). The mean values are calculated from four independent pull down experiments and their standard error is indicated. Red columns: RAC/ROPs assigned to pollen tube growth; Green columns: RAC/ROPs assigned to ABA response pathway. **(C)** Yeast transformants co-expressing either Nt-RAC5, Nt-RAC3, or Nt-RAC1 fused to the DNA binding domain of the GAL4 transcription factor (GAL4-DBD) together with Nt-RIC11pt fused to the GAL4 activation domain (GAL4-AD). Cells from yeast transformant single colonies were diluted in water and equal amounts plated on histidine-free culture medium (-His) to analyse for two-hybrid interaction. Equal amounts of all transformants were also plated on histidine containing medium (+His) as positive growth control. Controls for specific protein interaction were transformants co-expressing the Nt-RIC11pt prey protein with just the GAL4-DBD, as well as transformants co-expressing Nt-RAC5, Nt-RAC3, or Nt-RAC1 bait proteins together with just the GAL4-AD. **(D)** Phylogenetic analysis presented as unrooted radial tree obtained by maximum likelihood estimation and JTT matrix-based model providing protein amino acid sequence relationships between known ROPs of *Arabidopsis thaliana*, and RAC1, RAC3, RAC5, as well as RAC5.2 of *Nicotiana tabacum*, which were analysed in pull down experiment from (A) and yeast two hybrid assay from (C). The phylogeny reconstruction was generated using ‘MEGA-X 10.0.5 - software’. The indicated bootstrap values are based on 500 replications and the scale bar for branch lengths indicates the number of amino acid substitutions per site. At: *Arabidopsis thaliana*; Nt: *Nicotiana tabacum*; Red: RAC/ROPs assigned to pollen tube growth; Green: RAC/ROPs assigned to ABA response pathway.

In Y2H-assays Nt-RIC11pt fused to the activation domain of GAL4 (GAL4-AD:Nt-RIC11pt) interacted approximately equally with Nt-RAC1, Nt-RAC3, and Nt-RAC5 fused to the GAL4 DNA-binding domain (GAL4-DBD), whereby growth of yeasts demonstrating the interaction with Nt-RAC3 was slightly more pronounced (**Fig.5C**).

Taken together, these results suggest that tobacco Nt-RIC11pt is a versatile effector, which has the capacity to interact with multiple types of RAC-GTPases, what consequently implies association of Nt-RIC11pt with various cellular pathways. In particular, this suggests involvement in ABA-signalling via Nt-RAC3, and in pollen tube growth via Nt-RAC1, Nt-RAC5, and Nt-RAC5.2.

### 3.6 Nt-RIC11pt interacts with Nt-CAR4

To gain insights which signalling pathways are related to Nt-RIC11pt a Y2H-screen was performed particularly aiming to identify downstream targets using Nt-RIC11pt as bait. In this screen of a pollen tube library several cDNAs were identified comprising uncharacterized proteins of unknown function, and furthermore, a particularly interesting cDNA which encodes a 123 amino acids fragment of a protein belonging to a family of C2 (protein kinase C conserved region 2) calcium-dependent lipid-binding domain (CaLB domain) containing proteins (**Fig.6A**). BLAST search revealed that this protein is identical with the computer predicted sequence of tobacco Nt-CAR4/GAP1 (XP_016443372) and most closely related to the <u>G</u>TPase-<u>A</u>ctivating <u>P</u>roteins GAP1/CAR4 from Arabidopsis (AT3G17980; 76.4% identity) (**Fig.6A**), and Os-GAP1 from *Oryza sativa* (LOC4329192; 55.7% identity) (**Fig.6A**). These unconventional GAP-proteins contain a phospholipid-binding C2 domain and have been described to play a role in abscisic acid (ABA)-signalling, plant stress response, as well as TGN-related endomembrane trafficking (Cheung et al., 2013; Diaz et al., 2016; Heo et al., 2005). The closest human homolog is the *Homo sapiens* C2CD5 protein (C2 domain-containing protein 5) (EAW96476).

**Figure 6.**
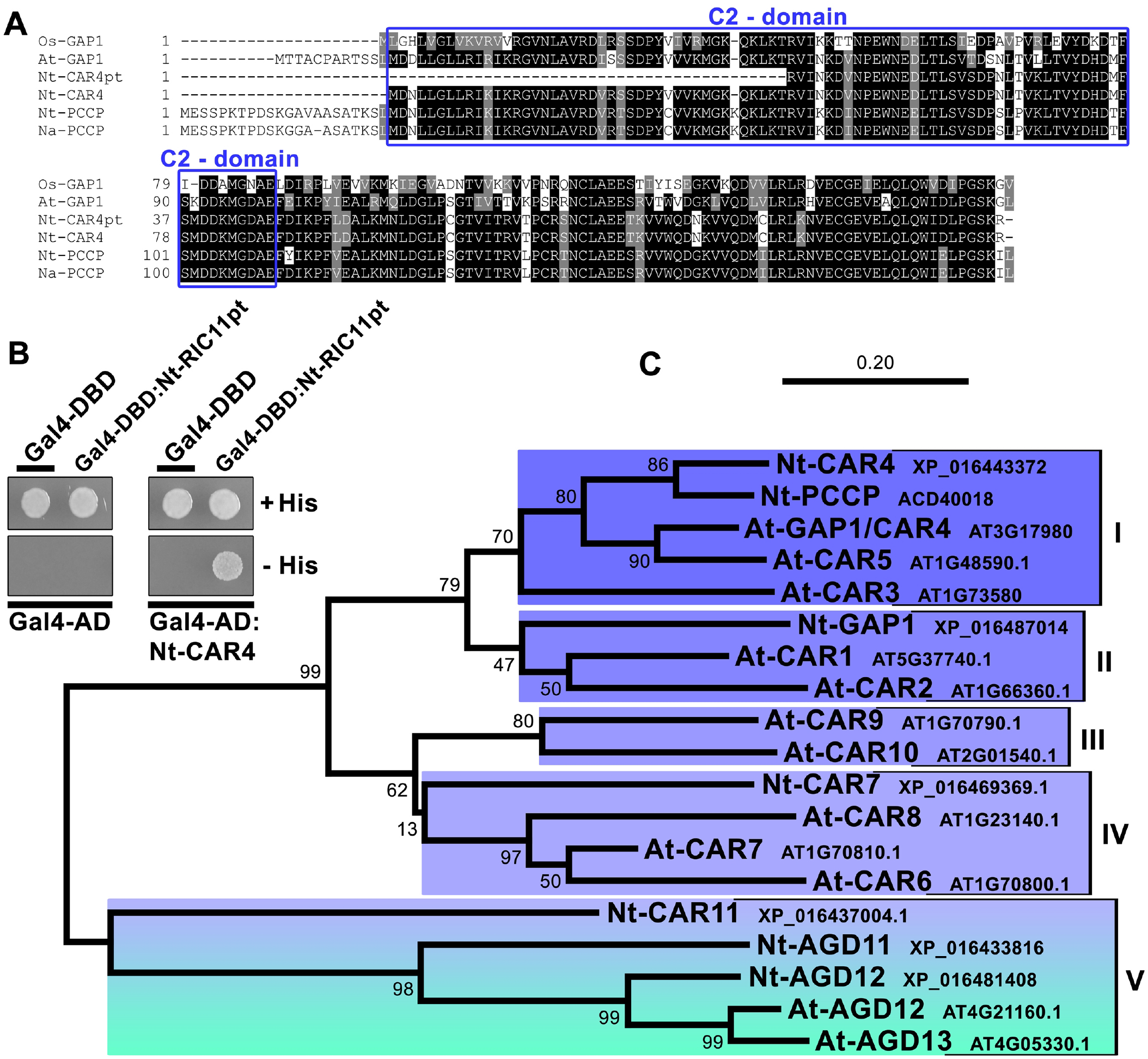
Yeast Two-Hybrid Assay, Amino Acid Sequence Alignment and Phylogenetic Analysis of the Nt-RIC11pt Interaction Partner Nt-CAR4. **(A)** Clustal Omega Amino Acid Sequence Alignment of Nt-CAR4 with closest homologs from *Arabidopsis thaliana (At), Nicotiana tabacum (Nt), Nicotiana alata (Na)*, and *Oryza sativa (Os)* identified by BLAST search. Black shading: sequence identity; Gray shading: sequence similarity; Blue box: C2-domain. **(B)** Yeast transformants co-expressing Nt-RIC11pt fused to the DNA binding domain of the GAL4 transcription factor (GAL4-DBD) together with Nt-CAR4 fused to the GAL4 activation domain (GAL4-AD). Yeast single colonies were diluted in water and equal amounts of transformed cells plated on histidine-free culture medium (- His) to analyse for two-hybrid interaction. All transformants were also plated on histidine containing medium (+His) as positive growth control. Controls for specificity of the protein interaction were transformants co-expressing the Nt-CAR4 prey protein with just the GAL4-DBD, as well as transformants co-expressing Nt-RIC11pt bait protein together with just the GAL4-AD. **(C)** Phylogenetic analysis delineated in an unrooted tree obtained by maximum likelihood estimation and JTT matrix-based model providing protein amino acid sequence relationships between Nt-CAR4 and closest homologs from *Arabidopsis thaliana* and *Nicotiana tabacum*. All members of the CAR-family were classified in V groups (various blue tones) according to their sequence similarity. Notice the remote clustering of Nt-CAR11 (blue) together with the ARF-GAPs AGD11-13 (green) in group V (blue to green shading). Phylogeny reconstruction was generated using ‘MEGA-X 10.0.5 - software’. The indicated bootstrap values are based on 500 replications and the scale bar for branch lengths indicates the number of amino acid substitutions per site. Locus/Accession number is given for every protein. At: *Arabidopsis thaliana*; Nt: *Nicotiana tabacum*.

Y2H-assays corroborated the result from cDNA library screening and provide clear evidence that Nt-CAR4 fused to the activation domain of GAL4 (GAL4-AD:Nt-CAR4) interacted specifically with Nt-RIC11pt fused to the GAL4 DNA-binding domain (GAL4-DBD) (**Fig.6B**).

Phylogenetic analysis of the most closely related Arabidopsis and predicted tobacco proteins reveals an unrooted tree of CAR/GAPs clustering in in five groups (**Fig.6C**). Nt-CAR4 aggregates with Nt-PCCP and At-GAP1/CAR4 in group I including At-CAR3 and At-CAR5 (**Fig.6C**). Furthermore, the tight clustering of Nt-CAR11 with AGD11, 12, and 13 clearly demonstrates that CAR/GAPs are closely related to ARF-GAPs (**Fig.6C**). Remarkably, Nt-CAR4 is most closely related to the pollen-specific C2-domain containing protein *Nicotiana alata* Na-PCCP (ACD40010; 85.3% identity) (**Fig.6A**) and the predicted homolog from *Nicotiana tabacum* Nt-PCCP (ACD40018; 84.5% identity) (**Fig.6C**).

The multifaceted CAR/GAP-family has the capacity for activating their interacting GTPases, therefore it is particularly interesting to analyse their relation to other known types of GAPs. For phylogenetic analysis documented sequences of the various Arabidopsis GTPase Activating Proteins from database were compiled and classified into three accepted families according to their known or predicted functions (Arf-GAPs, Rho/ROP-GAPs, Ypt/Rab-GAPs) and were complemented with Arabidopsis as well as tobacco CAR/GAPs. A phylogenetic tree was constructed using Maximum Likelihood estimation method with 500 bootstrap replications (**Fig.7**). This phylogenetic tree was generated from amino acid comparison and shows that members of the CAR/GAP family most tightly cluster and are additionally closely related to the broad distributed family of Arf-GAPs (ADP-ribosylation factor - GTPase-activating proteins), which form separate dispersed clusters (**Fig.7**). In particular, Nt-CAR11 is positioned near to Nt-AGD11/12 and At-AGD12/13 what indicates close relation of CAR/GAPs to the Arf-GAP family (**Fig.6C; 7**). In the displayed tree ARF-GAPs represent the most loosely clustered GAP-family with At-AGD14 and At-AGD11 being obviously very distantly related, even adjoining the families of Rho/ROP-GAPs or Ypt/Rab-GAPs (**Fig.7**).

**Figure 7.**
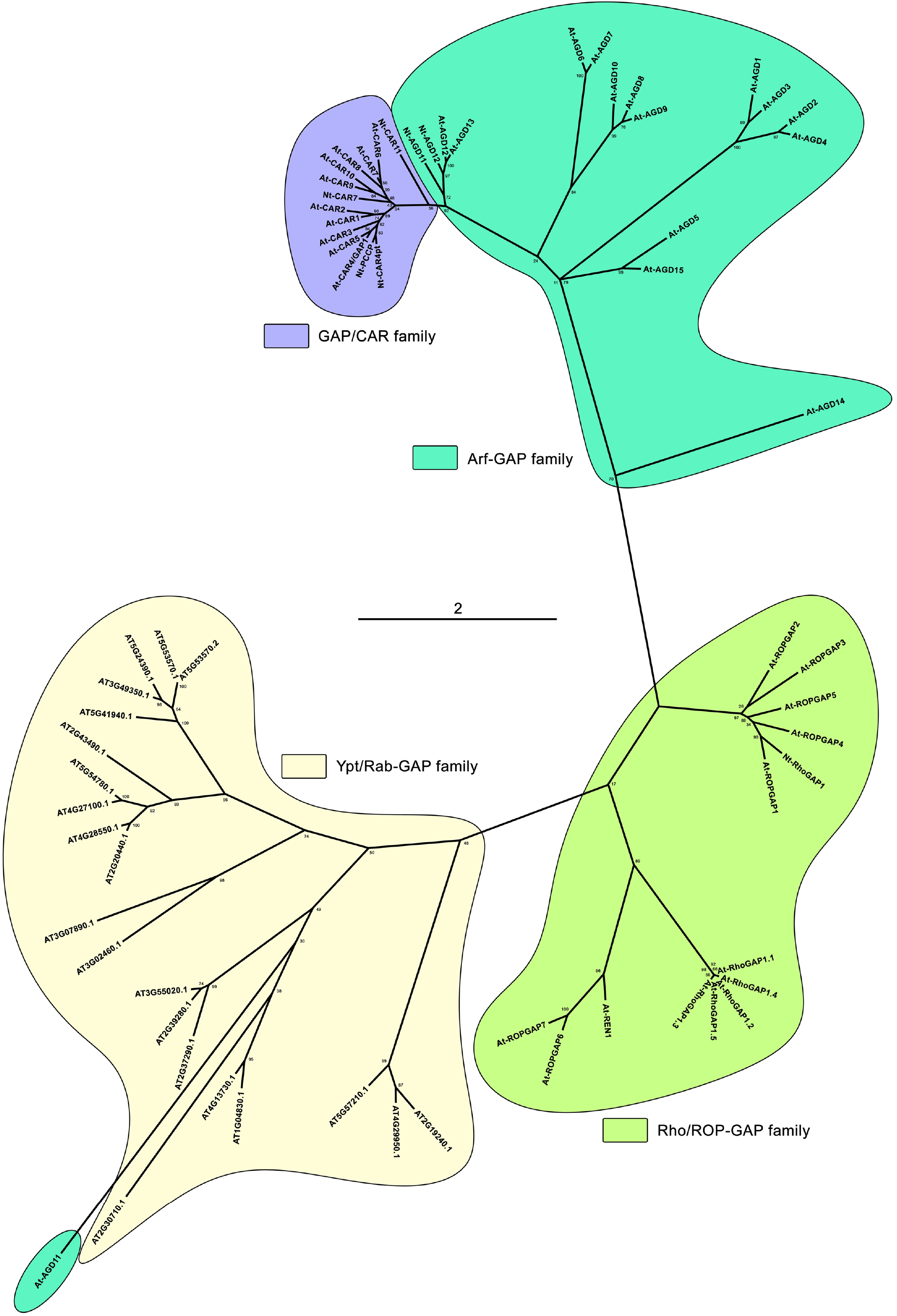
Phylogenetic Comparison of the CAR/GAP-family with known Arf-GAPs, Rho/ROP-GAPs and Ypt/Rab-GAPs. Phylogenetic analysis delineated in an unrooted tree obtained by maximum likelihood estimation and JTT matrix-based model providing protein amino acid sequence relationships between the CAR/GAP-family and known Arf-GAPs, Rho/ROP-GAPs and Ypt/Rab-GAPs. The CAR/GAP-family comprises known *Arabidopsis* *thaliana* proteins, Nt-CAR4pt, and computer predictions of *Nicotiana tabacum* from database search. Phylogeny reconstruction was generated using ‘MEGA-X 10.0.5 - software’. The indicated bootstrap values are based on 500 replications and the scale bar for branch lengths indicates the number of amino acid substitutions per site. Locus/Accession number is given for every protein. At: *Arabidopsis thaliana*; Nt: *Nicotiana tabacum*.

## 4. DISCUSSION

Plant pollen tubes are highly specialized cell types that reach outstanding growth velocities in order to transport the male gametes through the pistil to the ovule, therefore it is plausible that this remarkable cell type evolved pollen-specific molecular signal transduction networks to coordinate the cellular processes essential for traversing the female tissue.

To date, several predominantly pollen tube growth related regulatory proteins have been identified that belong to various functional categories, such as:

I) RAC/ROP-GTPases: At-ROP1 (Li et al., 1998) and Nt-RAC5 (Klahre and Kost, 2006),

II) RAC/ROP-effectors which are associated with actin- and membrane-dynamics: At-RIC3 (Gu et al., 2005; Lee et al., 2008b), Nt-ADF1 (Chen et al., 2003; Chen et al., 2002), Nt-RISAP (Stephan et al., 2014),

III) Receptor-like kinases (RLKs): At-ANX1/2 (Boisson-Dernier et al., 2013; Boisson-Dernier et al., 2009), At-MDIS1-MIK1/2 (Wang et al., 2016), At-PRK6 (Takeuchi and Higashiyama, 2016),

IV) The putative GTP-binding protein At-GPR1 (Yang et al., 2017), and V) Zm-CPK32 (Li et al., 2018), which is a calcium-dependent protein kinase that interestingly has been related to regulation of ABA-signalling.

In addition to these known proteins, the present work reveals that the RAC-effector Nt-RIC11pt belongs to such a preferentially in pollen expressed repertory of signal transduction molecules regulating pollen tube growth. In their effort to isolate all Arabidopsis RIC genes by PCR from cDNA libraries of flowers, seedlings, and leaves, Wu *et al*. (Wu et al., 2001) state that they could not amplify At-RIC11, what corroborates the observation that tobacco Nt-RIC11pt is exclusively expressed at high levels in pollen and pollen tubes. However, it cannot be ruled out that Nt-RIC11pt might also be expressed in other tissues at minor levels below the detection limit of utilized qRT-PCR.

The presented phylogenetic analyses show that Arabidopsis and tobacco RICs cluster together in 5 groups corresponding to the ones described by Wu *et al*. (Wu et al., 2001), whereby Nt-RIC11pt is categorized herein as a member of group I RICs (**Fig.2C**), whose functions have not been characterized to date.

### 4.1 Multifaceted RIC-RAC binding

It should be highlighted that data submitted in this work provide first evidence for interactions between RIC11 and different active RAC/ROP-GTPases from diverse phylogenetic groups performing individual functions. In particular, the unrooted tree in **figure 5D** shows that Nt-RAC3 clusters with At-ROP11 and At-ROP10 which both fulfil functions in ABA-signalling (Li et al., 2012a; Zheng et al., 2002). Whereas Nt-RAC1, Nt-RAC5, NtRAC5.2 group together with At-ROP1, At-ROP3, and At-ROP5, which have a role in actin dynamics and endomembrane traffic thus contributing to pollen tube growth (Feng et al., 2016; Gu et al., 2005; Lee et al., 2008b; Nibau et al., 2006). The specific involvement in these pollen tube growth related processes has been experimentally demonstrated for Nt-RAC1 (Chen et al., 2003) and Nt-RAC5 (Klahre et al., 2006; Stephan et al., 2014).

This allows the intriguing interpretation that RIC11 is a versatile effector representing a connective network hub which is involved in signal transduction to various plant pathways. Reports presenting data for other RICs support these findings, as they show that At-ROP1 binds to a range of 7 RICs (Wu et al., 2001), concluding that At-ROP1 thereby coordinates even antagonistic cellular processes, as for instance, in the case of RIC3 and RIC4 which are counteracting in actin dynamics during polarized tip growth of pollen tubes (Gu et al., 2005; Lee et al., 2008b). Additionally, RIC1 has been described to be involved in two different processes, namely actin filament severing in the pollen tube tip (Zhou et al., 2015), as well as microtubule organization in pavement cells (Fu et al., 2005).

The presented data in combination with observations of Wu *et al*. (Wu et al., 2001) and above mentioned studies allow to hypothesize that presumably every RIC might interact with every ROP. In this context, it is noteworthy that Arabidopsis expresses 11 ROPs and an equal number of RICs. If, by implication, RICs bind comprehensively to all ROPs this would constitute an interconnected RIC-ROP network, for which their highly conserved CRIB-domain might be the central binding-element. In this case the important question arises of how respective pathway-specificity is particularly attained additionally to the general CRIB-based interaction? There are several answers to this intriguing question.

Based on the high degree of conservation between CRIB-domains each RIC-family member holds the potential to bind all ROPs, however, heterogeneous flanking sequences adjacent to the CRIB-domain likely mediate differentiated interaction to individual ROPs (Thompson et al., 1998). The sequence alignments show that Nt-RIC11pt has several candidate sequences for this purpose. The high sequence diversity among RIC-family members outside of their highly conserved CRIB domain suggests the existence of various structural features potentially mediating differential interaction to different RACs or downstream targets. Most particularly, presented pull down data clearly demonstrate differential RIC-RAC binding, because Nt-RIC11pt interacts equally strong with RAC1, RAC5, and RAC5.2, whereas the affinity to RAC3 is approximately two fold higher. This might be caused by differential interacting domains, although it remains unclear which residues mediate the selective binding capacity of Nt-RIC11pt to diverse RACs.

In this context, the small and also highly conserved RAC-sequences must be diverse enough to contain elements mediating RIC specificity. It has to be highlighted, that various distinct regions outside of switch regions I and II in RAC-GTPases are known to be required to make contact to effector proteins (Bishop and Hall, 2000) and thus could potentially contribute to differential RIC-RAC binding.

Another potential specificity determinant might be post-translational modification of Nt-RIC11pt, for example combinatorial phosphorylation of its exceptional multitudinous serine/threonine residues, possibly through calcium-dependent protein kinases (CDPKs) which have been shown to be involved in abscisic acid (ABA)-mediated physiological processes (Mori et al., 2006; Zhu et al., 2007).

Furthermore, additional interaction partners might mediate pathway specificity, however, the presented *in vitro* pull down experiments verify a basic difference in the direct interaction between the various purified RACs and RIC11.

Most importantly, the *in vivo* localization analysis substantiated a universal cytoplasmic distribution of RIC11 what represents the essential characteristic for interaction with a wide-range of RACs. Consequently, the individual RAC-localization would determine the area of action for RIC11 and thus the affected pathway. In this case, simply the homogeneous cellular abundance of RIC11 would collectively involve various downstream events, what hence depicts RIC11 as interface for multiple RAC-dependent processes.

Besides that, the universal binding of RIC to different RACs could simply be interpreted as a backup system in which RICs are functionally redundant and potentially replace each other. However, this requires versatile binding not only to different GTPases but even to the diverse downstream targets. This would depict RICs as promiscuous and highly unspecific effectors, raising the issue why there are so many different types of RICs in one cell at all?

In any case, to fulfil its role as a universal switch of functionally diverse RACs, RIC11 would be toggled to the respective pathway and subsequently serve as a mediator to specific downstream regulators, such as CAR4.

### 4.2 What is the significance of CAR4-RIC11 interaction?

Nt-CAR4/GAP1 belongs to a poorly characterized family of ten C2-domain ABA-related (CAR) proteins which have been described to bind phospholipid membranes in a Ca^2+^-dependent manner and play roles in signal transduction based upon additional catalytic-activities, such as GAP-activity, kinase-activity, and mediation of protein-protein interactions (Diaz et al., 2016). In particular, members of the CAR-family recruit the pyrabactin resistance 1/PYR1-like (PYR/PYL) ABA-receptors to the membrane (Diaz et al., 2016; Rodriguez et al., 2014). The finding of Nt-RIC11pt interacting with ABA-related RAC3 and additionally Nt-CAR4/GAP1, which is associated with the PYR/PYL-receptor, indicates a role in the ABA-receptor pathway in pollen tubes.

The homologs of Nt-CAR4/GAP1, namely At-GAP1 and Os-GAP1 show both GTPase/ATPase activities and phospholipid binding capacities (Cheung et al., 2013; Cheung et al., 2010). Interestingly, the presented phylogenetic analyses demonstrate that CAR/GAPs tightly cluster as a distinct lateral arm closely linked to the broad family of diverse structured Arf-GAPs, which analogously to CAR/GAPs are multifunctional in addition to their Arf-GAP activity, and have domains that differ from those of other GAPs. In contrast to classical RhoGAPs, such as Nt-RhoGAP1 (Klahre and Kost, 2006), the protein sequences of CAR/GAPs apparently contain no intrinsic CRIB-domain, which in classical RhoGAPs is responsible for ROP-binding and required for efficient GAP activity (Klahre and Kost, 2006; Schaefer et al., 2011; Wu et al., 2000). Therefore the present work suggests that the effector RIC11 fulfils the function of recruiting CAR4/GAP1 to the respective RAC/ROP-GTPase.

Os-GAP1 and At-GAP1 have been described to be involved in plant stress and defence response by stimulating GTPase-activity of the unconventional G-protein YchF1, which belongs to the poorly characterized YchF-type family with largely unknown functions (Cheung et al., 2013; Cheung et al., 2008). Noteworthy is that Os-GAP1 has been connected to TGN-related endomembrane transport by regulating Os-Rab8a and Os-Rab11 (Heo et al., 2005; Son et al., 2013).

Additionally, the CAR4-homolog Na-PCCP is a pollen-specific protein that associates with signal proteins from extracellular matrix (ECM) of the pistil (Lee et al., 2008a). The C-terminal region of Na-PCCP binds to arabinogalactan proteins which are endocytosed from ECM of the pistil and have a function in promoting pollen tube growth by providing a chemical guidance signal (Lee et al., 2009; Lee et al., 2008a). This allows to hypothesize that the RIC11-CAR4 interaction might contribute to pistil-pollen crosstalk, and moreover, implicates a potential role in endomembrane transport, because the C2-domain of Na-PCCP associates with the plasma membrane and the endosomal system (Lee et al., 2009).

CAR proteins are partially cytosolic localized calcium sensors with basic membrane association that upon increase of Ca^2+^-level is enhanced through structural rearrangement of the cell membrane induced by CARs (Diaz et al., 2016). Furthermore, it has been proposed that CARs might act as a scaffold proteins that recruit interaction partners like PYR/PYL-receptors to signalling platforms in specific plasma membrane domains (Diaz et al., 2016). Thus, a minor fraction of the cytoplasmic RIC11-population might be transiently associated with subdomains of the plasma membrane via interaction with CAR4 and membrane bound RAC3. Correspondingly, the function of At-ROP10 and At-ROP11 in PYL/PYR-related ABA-signalling is modulated by intracellular calcium levels (Li et al., 2012a; Mori et al., 2006; Zhu et al., 2007). In this context, the tip-directed calcium gradient of pollen tubes, which is a prerequisite for elongation (Steinhorst and Kudla, 2013), potentially causes an equivalent aligned accumulation of the calcium sensor CAR4 in the plasma membrane, where RIC11 could establish spatial contact to membrane-bound RACs. However, an apical accumulation of CAR4 in the tube plasma membrane, and the subcellular interaction site with RIC11 remains to be substantiated.

### 4.3 Which cellular processes are targeted by RIC11?

In this context binding to RAC1 and RAC5 implies a role in endomembrane transport and actin-remodelling during polarized pollen tube growth (Chen et al., 2003; Kost, 2008; Stephan, 2017). It has to be highlighted that Nt-RIC11 most significantly interacts with Nt-RAC3 which is the tobacco homolog of At-ROP10, and additionally, Nt-RAC3 phylogenetically closely clusters with At-ROP9 and At-ROP11 (**Fig.5D**), what synoptically suggests a role in ABA-signalling of pollen tubes. Interestingly, the few non plant homologs of RIC11 are either related to rearrangement of the actin cytoskeleton as the Cdc42-effector WASP from *Homo sapiens* (Kim et al., 2000), or are particularly involved in pheromone response and pseudohyphal/invasive growth as *Saccharomyces cerevisiae* STE20 (Leberer et al., 1992).

In this context, several studies show that ROPs are important factors in cellular response to plant hormones, in particular Arabidopsis ROP2, ROP6, ROP9, ROP10, and ROP11 are related with ABA-signalling (**Fig.5D** green), affecting seed germination, root elongation, stomatal closure, and embryo development (Lemichez et al., 2001; Li et al., 2001; Li et al., 2012a; Nibau et al., 2013; Zheng et al., 2002). Most interestingly, transgenic expression of constitutive active ROP10 in Arabidopsis reduces ABA-response and consistently a rop10 null mutant is hypersensitive to ABA (Zheng et al., 2002). Similar results for At-ROP11 (Li et al., 2012a) show that it acts in parallel with At-ROP10 as a negative regulator of multiple ABA-responses. In contrast, At-ROP9 has been reported to act antagonistically to ROP10 and ROP11 in abscisic acid signalling during embryo development and lateral root formation (Nibau et al., 2013).

In this regard, it is also worth noting that all cluster members, namely ROP9, ROP10, and ROP11 belong to a subgroup of RAC/ROP-GTPases, which specifically evolved only in vascular plants (Winge et al., 2000), hence implicating that Nt-RIC11 gained its function with the emergence of complex land plants, what in turn corresponds to the observation that Nt-RIC11 is expressed pollen-specific.

Directional pollen tube growth is susceptible to molecular factors that are exchanged between the female pistil and the pollen tube, such as small secreted proteins, which play a role in pollen tube adhesion and guidance (Chae and Lord, 2011). In addition, other exogenous molecules, as the phytohormone ABA might potentially influence tube elongation *in vivo*, however, a related role for ABA has not been studied in depth. Some studies provide evidence, that according to its concentration ABA effects positive and negative changes in growth velocity of the male gametophyte (Dhawan and Malik, 1981; Frascaroli and Tuberosa, 1993; Malik and Chhabra, 1976; Wu et al., 2008; Yang et al., 2003). Low exogenous ABA levels promote tube growth what correlates with falling ABA concentration in the female tissue upon flowering and pollination (Kojima et al., 1993). In accordance therewith, ROP10 gates the expression of genes that are specific to low concentrations of ABA (Xin et al., 2005). Furthermore, Kovaleva *et al*. (Kovaleva et al., 2016) show that ABA-induced lateral redistribution of H^+^-ATPase into the subapical zone of pollen tubes and increased the content of microfilaments accompanied by a redistribution of F-actin to apical zone. This demonstrates a role for ABA in remodelling of the subcellular tip-composition.

Taken together, this study suggests that Nt-RIC11pt acts as a potential network hub between Nt-RAC1-, Nt-RAC5-, and Nt-RAC3-related pathways, thus coordinating actin-dynamics and endomembrane transport with ABA-signalling during pollen tube growth (**Fig.8**). In this context it is hypothesized that ABA-response in tobacco pollen tubes may be positively affected through deactivation of the ROP10-homolog Nt-RAC3 via recruitment of Nt-CAR4/GAP1 by Nt-RIC11pt (**Fig.8**). In concordance therewith, it has been shown that Feronia suppresses ABA-response via GEFs that activate ROP10, which in turn, activates ABI2 (Yu et al., 2012; Zhao et al., 2015).

It has to be mentioned that only few functions have been specified to date for any of the Arabidopsis RICs that phylogenetically cluster in group I (**Fig.2C**). Interestingly, Wu *et al*. (Wu et al., 2001) state that RIC10 promotes pollen tube growth. Furthermore, concerning the involvement of other RICs in ABA-signalling some data exist. For instance, LLP-12-2 from *Lilium longiflorum*, which is a homolog of At-RIC6 belongs to RICs of group III (Data not shown) (Hsu et al., 2010) and has been identified as effector of LLP-ROP1, an At-ROP1 homolog (Hsu et al., 2010). Interestingly, in *Lilium* pollen tubes LLP-ROP1 and its effector LLP-12-2 antagonistically respond to exogenous ABA application (Hsu et al., 2010).

Additionally, the well-characterized At-RIC1, which is expressed in many Arabidopsis tissues, has been reported to suppress ABA-response in germinating seeds and growing roots (Choi et al., 2013). However, Choi *et al*. (Choi et al., 2013) do not assign any ROP to the ABA-related function of At-RIC1, but they additionally state that the observed moderate phenotype of their ric1 mutants might be explained by an involvement of other RICs in ABA-signal transduction.

### 4.4 Epilogue

Besides the reported counter-action of two RIC-effectors (Gu et al., 2005; Hwang et al., 2005; Wu et al., 2001), the present study provides an additional mechanism whereby RIC11 might join together both functions, a recruitment of active RAC to target complexes, as well as deactivation by relaying GAP-activity, thus mediating negative feedback (**Fig.8**). Therefore, the reported RAC-RIC11-CAR4 network provides another mode of RAC/ROP-GTPase control in addition to classical Rho-GAPs and guanine-nucleotide-dissociation inhibitors (GDIs).

Altogether, the current data allow to postulate RIC11 as a multifaceted scaffold with differential binding efficiency to RAC/ROPs, performing effector-functions by mediating signals to various downstream pathways, as well as regulator-functions by additionally relaying negative feedback to GTPases (**Fig.8**). Furthermore, its eclectic interaction capacity opens up the possibility that RIC11 is a network hub interconnecting cytoskeleton dynamics, membrane transport processes, and ABA-controlled cellular water balance, which altogether control pollen tube elongation most likely in a coordinated manner.

**Figure 8.**
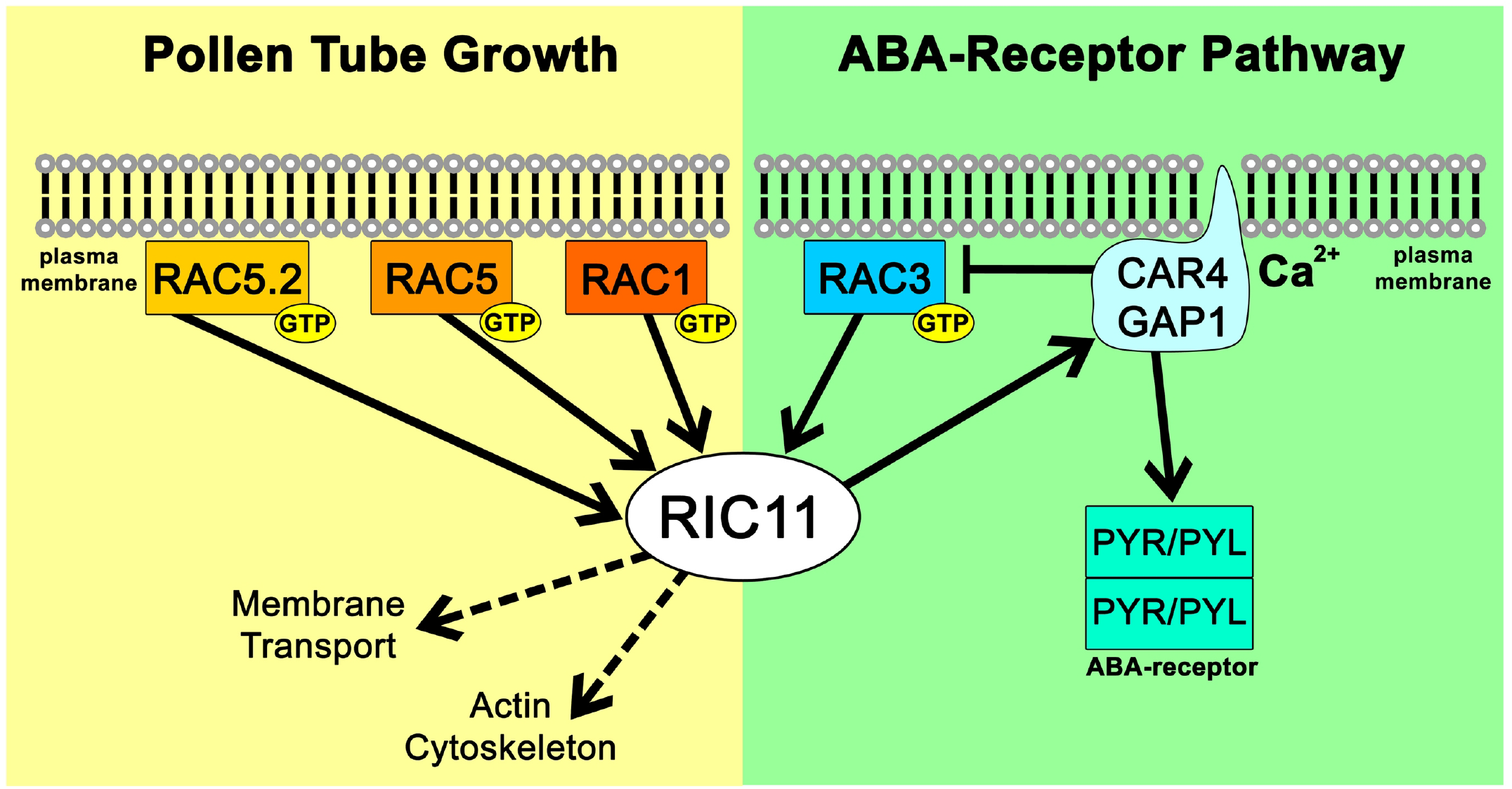
Graphical Overview of RIC11 Interaction Network. Schematic view of the RIC11 interaction network, including different RACs and CAR4/GAP1. The connection RAC3-RIC11-CAR4/GAP1 provides a link to the related PYR/PYL-homodimer, thus establishing a role in ABA-response. Cytoplasmic RIC11 indirectly mediates GAP-activity to membrane-associated RAC3 by recruitment of the unusual membrane-bound CAR4/GAP1. Ca^2+^-dependent membrane association of CAR4/GAP1 establishes a connection to the intracellular calcium gradient, which is a prerequisite for pollen tube growth. Solid arrows indicate steps supported by experimental data described in this paper and additional studies (Cheung et al., 2013; Cheung et al., 2008; Diaz et al., 2016; Rodriguez et al., 2014), whereas dotted arrows indicate implicated more putative relations based on other currently available data (Stephan et al., 2014).

## 5. ACKNOWLEDGEMENT

I would like to thank Hildegard Stephan for comments and discussions.

